# An interactome landscape of SARS-CoV-2 virus-human protein-protein interactions by protein sequence-based multi-label classifiers

**DOI:** 10.1101/2021.11.07.467640

**Authors:** Ho-Joon Lee

## Abstract

The new coronavirus species, SARS-CoV-2, caused an unprecedented global pandemic of COVID-19 disease since late December 2019. A comprehensive characterization of protein-protein interactions (PPIs) between SARS-CoV-2 and human cells is a key to understanding the infection and preventing the disease. Here we present a novel approach to predict virus-host PPIs by multi-label machine learning classifiers of random forests and XGBoost using amino acid composition profiles of virus and human proteins. Our models harness a large-scale database of Viruses.STRING with >80,000 virus-host PPIs along with evidence scores for multi-level evidence prediction, which is distinct from predicting binary interactions in previous studies. Our multi-label classifiers are based on 5 evidence levels binned from evidence scores. Our best model of XGBoost achieves 74% AUC and 68% accuracy on average in 10-fold cross validation. The most important amino acids are cysteine and histidine. In addition, our model predicts experimental PPIs with higher accuracy than text mining-based PPIs by 4% despite their smaller data size by more than 6-fold. We then predict evidence levels of ∼2,000 SARS-CoV-2 virus-human PPIs from public experimental proteomics data. Interactions with SARS-CoV-2 Nsp7b show high evidence. We also predict evidence levels of all pairwise PPIs of ∼550,000 between the SARS-CoV-2 and human proteomes to provide a draft virus-host interactome landscape for SARS-CoV-2 infection in humans in a comprehensive and unbiased way *in silico*. Most human proteins from 140 highest evidence predictions interact with SARS-CoV-2 Nsp7, Nsp1, and ORF14, with significant enrichment in the top 2 pathways of vascular smooth muscle contraction (CALD1, NPR2, CALML3) and Myc targets (CBX3, PES1). Our prediction also suggests that histone H2A components are targeted by multiple SARS-CoV-2 proteins.

## INTRODUCTION

The new coronavirus species of SARS-CoV-2 has been an unprecedented global threat causing COVID-19 disease with more than 5.6 million deaths over 2 years as of February 1, 2022 (https://covid19.who.int/). The scientific community has been working towards a better understanding of the biology and disease of SARS-CoV-2 infection and COVID-19 and a cure or a medicine is yet to be discovered, although several vaccines have been successfully developed and administered in many countries. In response to this global challenge, we previously studied network controllability of a human directed protein-protein interaction (PPI) network for SARS-CoV-2 infection using proteomics and other omics data (Lee, 2021). Here we aim to develop machine learning models to predict a degree of evidence or confidence for virus-human physical or functional PPIs for SARS-CoV-2 infection. Previous studies have focused on binary classification tasks of pathogen-host PPIs including SARS-CoV-2 (Alguwaizani et al., 2018; Cui et al., 2016; Dey et al., 2020; Ding and Kihara, 2018; Du et al., 2021; García-Pérez et al., 2018; Kshirsagar et al., 2021; Nourani et al., 2015, 2016; Sen et al., 2016; Yang et al., 2020; Zhang et al., 2017). Machine learning has been widely used in PPI prediction in general (Sarkar and Saha, 2019). Although binary classification for PPIs has been successful, measurements of physical PPIs are often noisy and subject to a rather subjective threshold for high or low confidence (Gordon et al., 2020a; Gordon et al., 2020b), which has been also studied in our previous work (Lee, 2021). In addition, such binary classifiers trained with data of physical PPIs are not directly generalizable to functional PPIs of many different types such as co-expression or common pathways.

To address the limitation of binary classification, we use large-scale virus-host PPI data with evidence scores for different types of PPIs from a public database, Viruses.STRING (Cook et al., 2018). We build multi-label classification models by binning evidence scores into 5 evidence classes or levels. To the best of our knowledge, no study has been done using the database for multi-label classification of virus-host PPIs. We use two tree-based classifiers in this study: random forests for bagging (Breiman, 2001) and XGBoost for boosting (Chen and Guestrin, 2016). While bagging algorithms only control for high variance in a model, boosting algorithms control both bias and variance and hence are considered to be more effective. In particular, XGBoost algorithms have shown superior performances in many different applications, as was the case in our previous study (Smith et al., 2022). As for model features, we compute a number of different similarity measures between amino acid composition profiles of virus-human interacting protein pairs. In other words, we construct protein sequence-based multi-label classifiers to predict virus-human PPIs with different evidence or confidence levels. Feature importance is examined by two alternative methods. We take the best models to apply to public experimental proteomics data of SARS-CoV-2 infection, which is not included in the training data of Viruses.STRING. In addition, we predict evidence levels for all PPI pairs between the SARS-CoV-2 and human proteomes in a comprehensive and unbiased way to provide a draft virus-host interactome landscape of SARS-CoV-2 infection in humans *in silico*. Finally, we prioritize those virus-human PPIs or sub-networks of high evidence for biological relevance and therapeutic opportunities for SARS-CoV-2 infection and COVID-19 in future studies.

## METHODS

### Data for model training and validation

We use the Viruses.STRING database (v10.5) for model training and validation. It contains 80,775 virus-human PPIs, excluding SARS-CoV-2. Each PPI has a combined score, ranging from 0 to 1000, combined from multiple scores of different PPI types such as physical experiments, co-expression, co-occurrence, homology, or text mining. We take combined scores as PPI evidence and bin the scores into 5 evidence classes (ECs) with bin size = 200 for multi-label classification: EC1 = 27,990 PPIs, EC2 = 33,001 PPIs, EC3 = 14,642, EC4 = 4,037, and EC5 = 1,105 PPIs. 6,684 PPIs are based on experiments with combined scores >= 435 (i.e., EC >= 3; 6,481 PPIs in EC3). On the other hand, 40,465 PPIs are from text mining with 89.4% PPIs having combined scores < 400 (20,447 PPIs in EC1 and 15,734 PPIs in EC2). Therefore, we pay particular attention to the two subsets of PPIs as part of prediction controls in this work.

### Test data

Our test data is 1,998 SARS-CoV-2 virus-human PPIs from the IntAct database as of July 17, 2020. We also tested all 549,990 protein pairs between the SARS-CoV-2 and human proteomes (27 SARS-CoV-2 proteins and 20,370 reviewed proteins from UniProt).

### Model features

For feature engineering, we first consider a virus-host undirected PPI network for generality, where nodes are virus or host proteins and edges are virus-host PPIs. In this work we are not concerned with PPIs among virus proteins or human proteins. In other words, one can think of it as an undirected bipartite graph. Given such a network or graph, we compute fractional compositions of the standard 20 amino acids of individual protein sequences as node features. We then compute various distance/similarity measures between the node features of each virus-host PPI pair as edge feature. We use the following 72 measures in this work: Pearson correlation, dot product, cross-correlation, Euclidean distance, cosine distance, Manhattan distance, Minkowski distance, Jensen-Shannon distance, Chebyshev distance, Canberra distance, Bray-Curtis distance, mutual information, the difference for each amino acid (human – virus), the absolute difference for each amino acid, and the ratio for each amino acid (virus/human). All computations were done in Python.

### Models

We use random forests (RF) and XGBoost (XGB) as multi-label classifiers (*sklearn* and *xgboost* packages in Python). As XGB model was the best performing classifier in our previous work (Smith et al., 2022) as well as in other studies (Chen and Guestrin, 2016), we chose XGB as our main model in this work and RF as a baseline. We trained 36 RF models by grid search and performed 10-fold cross validation (CV) for each model (i.e., a total of 360 fits) with the following parameter values: n_estimators = (200, 500, 1000); max_samples = (1.0, 0.75, 0.5); criterion = (gini, entropy); max_features = (sqrt, log2). We trained 432 XGB models by grid search and performed 10-fold CV for each model (i.e., a total of 4,320 fits) with the objective function of the soft probability and the following parameter values: n_estimators = (200, 500, 1000); max_depth = (3, 6); learning rate = (0.05, 0.1, 0.3); gamma = (0.0, 1.0); reg_lambda = (1.0, 2.0); reg_alpha = (0.0, 1.0); subsample = (1.0, 0.75, 0.5). For both RF and XGB models, the random seed was 1618 and the scoring was based on the weighted one-vs-rest AUC score and the accuracy. The refit was done by AUC. The split ratio between training and validation of the viruses.STRING data is 80:20 (or 64,620 and 16,155 PPIs, respectively).

### Analysis of feature importance

We used the impurity-based importance and the game-theoretic Shapley analysis to identify important features for prediction. Feature importance values are provided by the fitted classifiers in the *sklearn* and *xgboost* packages in Python based on mean decrease in impurity. For the Shapley analysis, we use the *shap* package in Python for analysis of SHAP (SHapley Additive exPlanations) values, as in our previous work (Smith et al., 2022).

### Functional analysis of human interacting proteins

We analyzed biological pathways enriched among human proteins interacting with SARS-CoV-2 proteins from our predictions. To do this, we used a webtool, *Enrichr* (https://maayanlab.cloud/Enrichr/) (Xie et al., 2021). We also performed protein-protein association analyses from the STRING database (STRING v11.5) (Szklarczyk et al., 2021). Network visualization was done using *Cytoscape* (Shannon et al., 2003).

## RESULTS

### Model performance

An overview of our modeling framework is shown in Fig. 1A. The mean cross-validated AUC by the best RF model, denoted *RF\**, is 67% with the following parameters: n_estimators = 500, max_samples = 0.75, criterion = *entropy*, and max_features = *sqrt*. The mean cross-validated AUC by the best XGB model, denoted *XGB\**, is 74% with the following parameters: gamma = 0.0, learning_rate = 0.3, max_depth = 6, n_estimators = 1000, reg_alpha = 1.0, reg_lambda = 1.0, and subsample = 1.0. Details of example performances for a 20% test set of 16,511 PPIs (random seed = 1618) by *RF\** and *XGB\** are given in Figs. 1B and 1C. The prediction accuracies for all 80,775 PPIs by *RF\** and *XGB\** are 92.0% and 93.5%, respectively.

**Figure 1.**
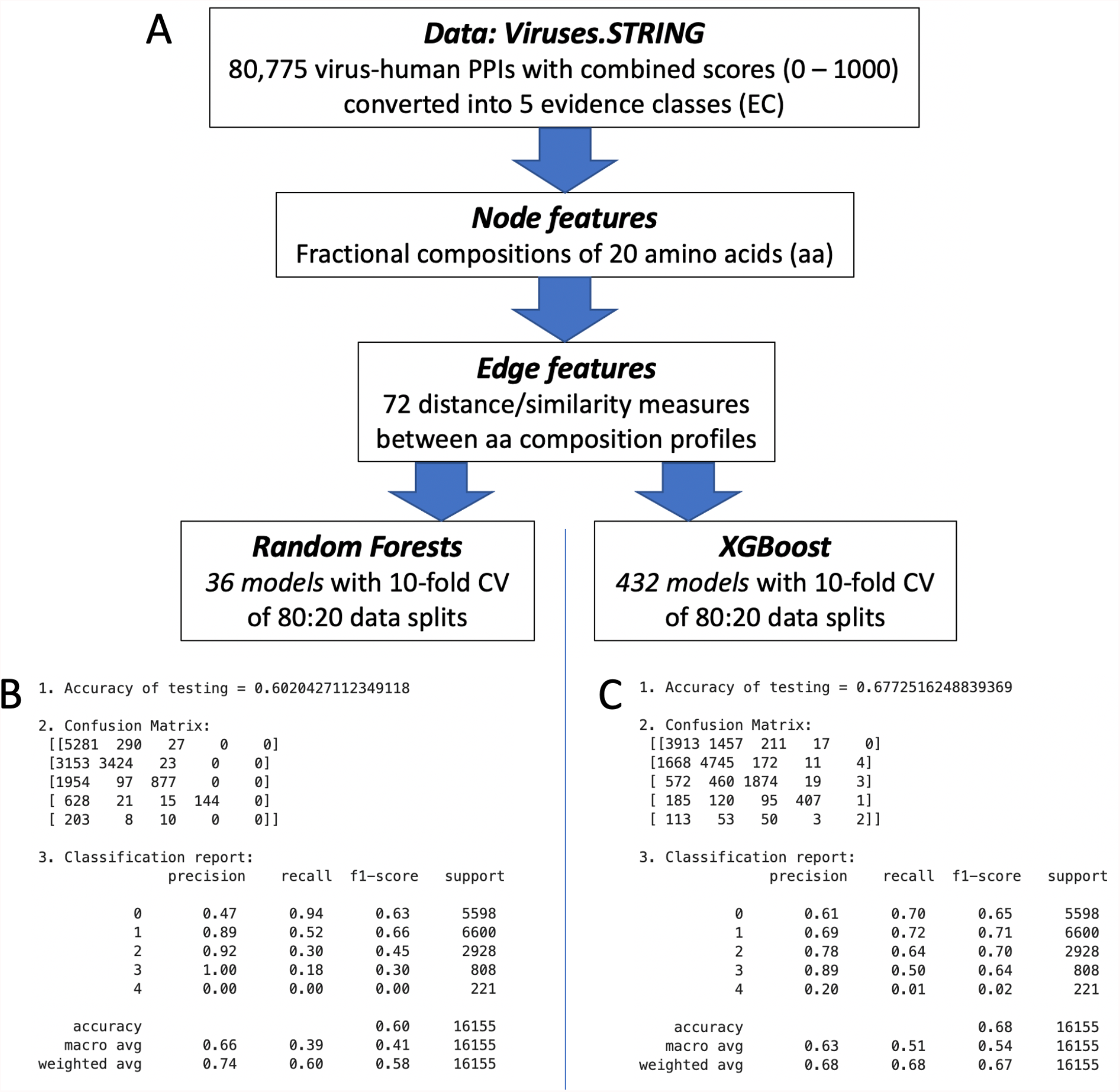
An overview of our modeling framework. (A) A flowchart of our pipeline. (B and C) Performance measures by *RF\** (B) and *XGB\** (C) for an example test set.

### Feature importance

As *XGB\** performed better than *RF\**, we focused on identifying important features for prediction by *XGB\**. We used both impurity-based importance and the SHAP analysis for all data. With impurity-based importance, the top 2 features are differences in cysteine and histidine fractions (C_minus and H_minus) (Fig. 2A). The SHAP analysis shows that ratios in cysteine and histidine fractions (C_ratio and H_ratio) are the top 2 features with the highest impact on model outputs on average. C_ratio and H_ratio have most impact on prediction of PPIs with EC3 and EC1, respectively (Figs. 2B and 2C). In particular, SHAP value distributions for each evidence class show that low C_ratio and low H_ratio have negative impact on EC3 and EC1 prediction for subsets of samples, respectively (Fig. 2C).

**Figure 2.**
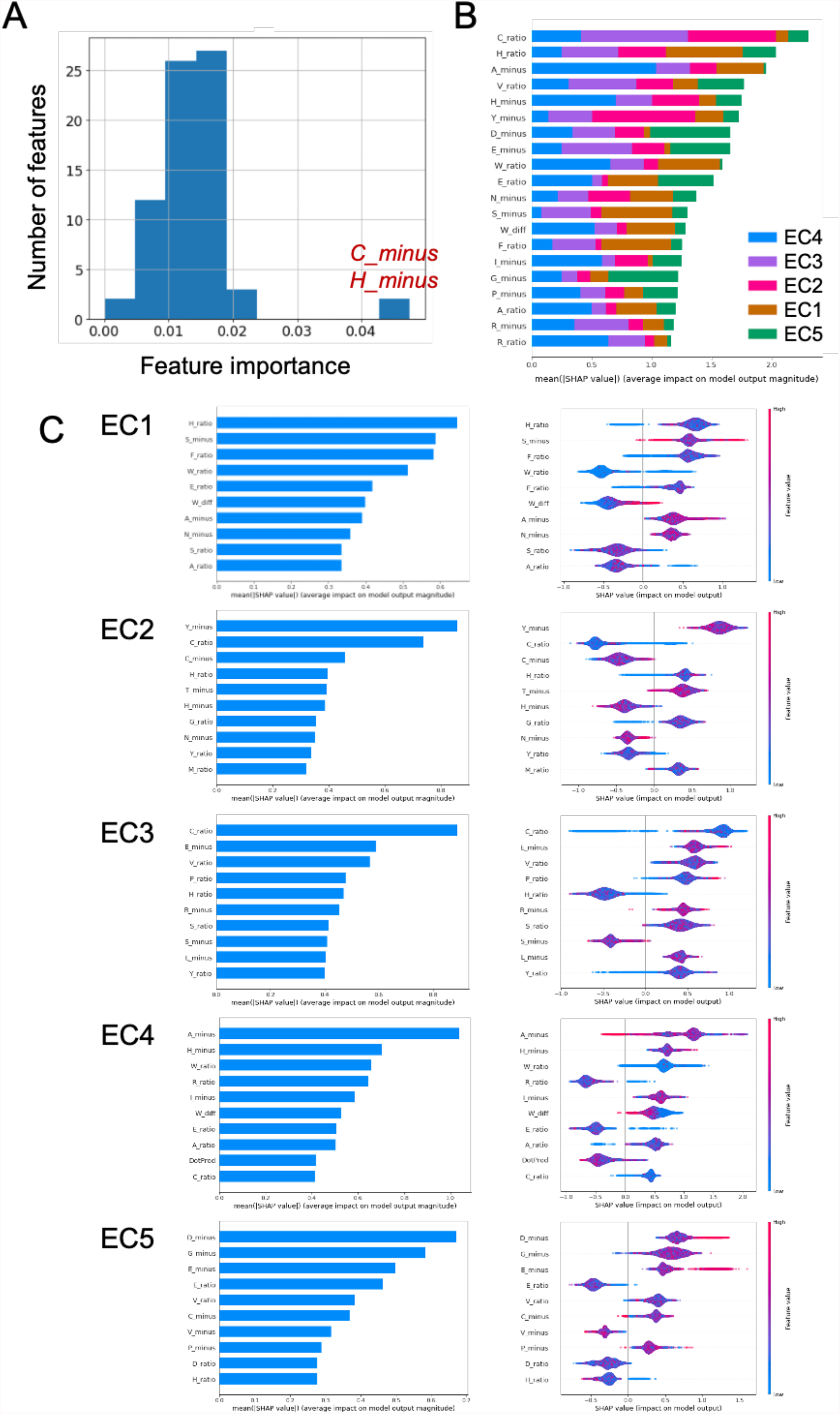
Analysis of feature importance. (A) A histogram of impurity-based feature importance values. The top 2 features, *C_minus* and *H_minus*, are indicated. (B) A bar plot of the means of absolute SHAP values for the top 20 features. (C) Average impact (bar plots) and individual impact (dot plots) of the top 10 features on model outputs for each evidence class by SHAP analysis.

### Prediction of experiments-based and text mining-based PPIs

As most experiments-based PPIs are in EC3 and most text mining-based PPIs are in EC1 and EC2, we applied *XGB\** and *RF\** to the two PPI subsets as a control of model performance. *XGB\** predicts 6,684 experimental PPIs with higher accuracy of 94% than 40,465 text-mining-based PPIs of 90%. On the other hand, *RF\** predicts the experimental PPIs with lower accuracy of 88% than the text mining-based PPIs with 91% accuracy. The largest number of predictions for the experimental PPIs occurred with EC3 for both *XGB\** and *RF\** (92.1% and 85.5%, respectively), as expected (Fig. 3A). The overlap of the EC3 predictions from both models is 85.4% (Fig. 3A). The largest number of predictions for the text mining-based PPIs occurred with EC1 for both *XGB\** and *RF\** (51.1% and 58.1%, respectively), and similarly for EC2, as expected (Fig. 3B). The overlap of the EC1 predictions from both models is 50.8% (Fig. 3B). We also performed a control experiment where we predicted 100 sets of random 6,684 text-mining-based PPIs (the same number as that of the experimental PPIs) using both models. The largest number of predictions remained to occur with EC1 (Fig. 3C). Therefore, despite the disadvantageous imbalance of experimental PPIs in the training data, predictions of EC3 are likely to suggest physical PPIs with experimental support.

**Figure 3.**
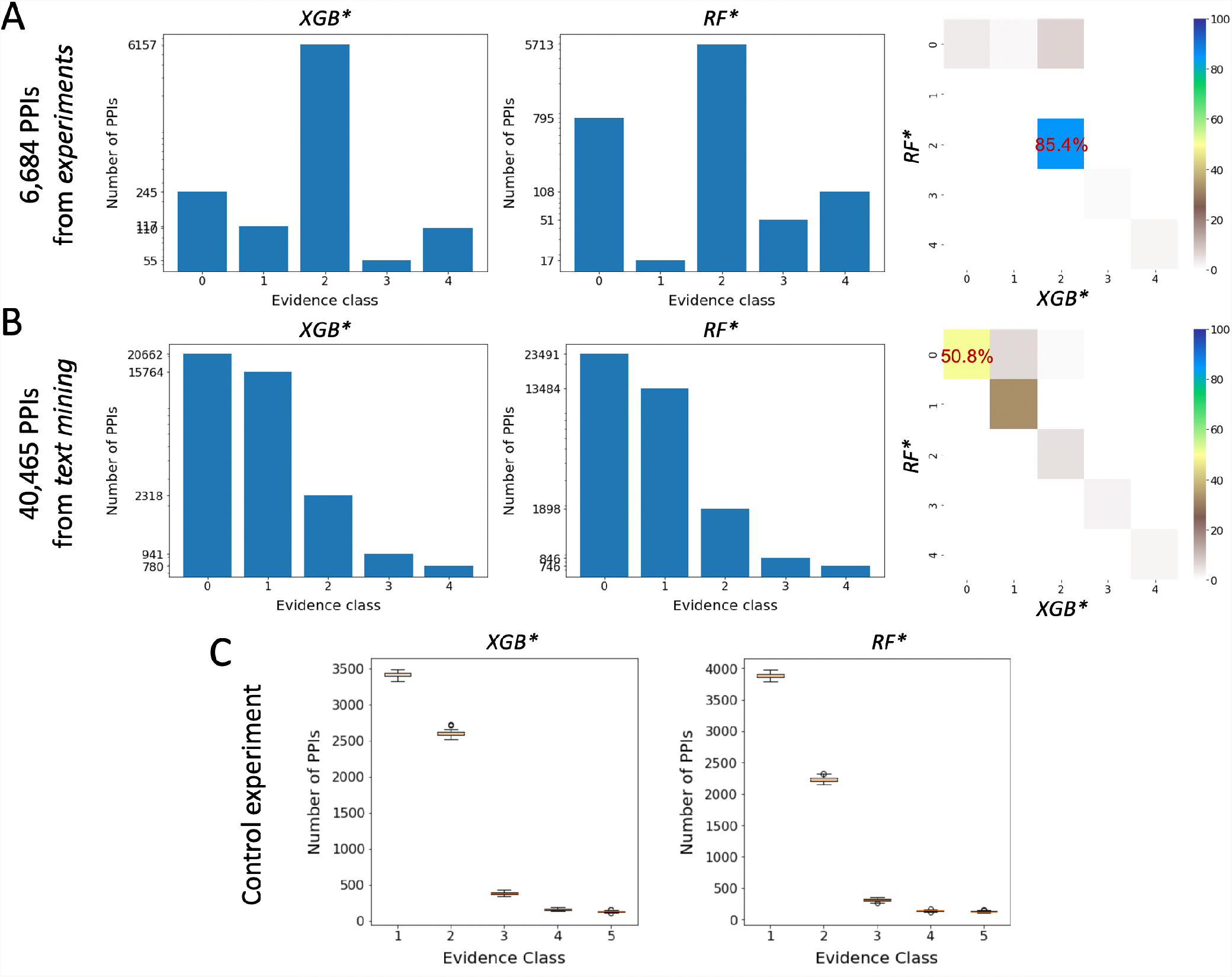
Model performance for experiments-based and text mining-based PPIs. (A and B) Prediction distributions for experiments-based PPIs (A) and text mining-based PPIs (B) by *XGB\** and *RF\**, and heat maps of prediction agreements between the two models. Note that the evidence class values are 0-index based. (C) A control experiment of predictions of random subsets of text mining-based PPIs whose size is the same as that of experiments-based PPIs.

### Prediction for SARS-CoV-2 virus-human PPIs

Given the model performance and evaluation above, we applied *RF\** and *XGB\** to SARS-CoV-2 experimental PPI data from the IntAct database. *RF\** predicted 1,916 PPIs as EC1, 26 PPIs as EC2, and 56 PPIs as EC3 (Table S1). *XGB\** predicted 1,072 PPIs as EC1, 426 PPIs as EC2, 487 PPIs as EC3, and 13 PPIs as EC4 (Table S1). Predictions of 1,142 PPIs agree between the RF and XGB models (Fig. 4A). In particular, 53 PPIs are predicted with EC3 by both models and all PPIs are involved with SARS-CoV-2 protein, Nsp7b. Those interacting human proteins are enriched in oxidative phosphorylation (ATP5F1A, ATP5F1B, and ATP6AP1) and SNARE interactions in vesicular transport (GOSR1, STX2, STX5, STX6, and VTI1A) (https://maayanlab.cloud/Enrichr/enrich?dataset=0120e7db0fb18ea9a357d7567c9b0003). Those 13 PPI predictions with EC4 predicted by *XGB\**, but not by *RF\**, are between 6 SARS-CoV-2 proteins and 13 human proteins. The 13 human proteins are enriched in cholesterol/steroid biosynthesis (MSMO1) and Myc targets (SLC25A3 and SSBP1) (https://maayanlab.cloud/Enrichr/enrich?dataset=02c0c07b4eec715852411f6a93bc90d9). The prediction differences can be easily seen in network visualization (Figs. 4B and 4C).

**Figure 4.**
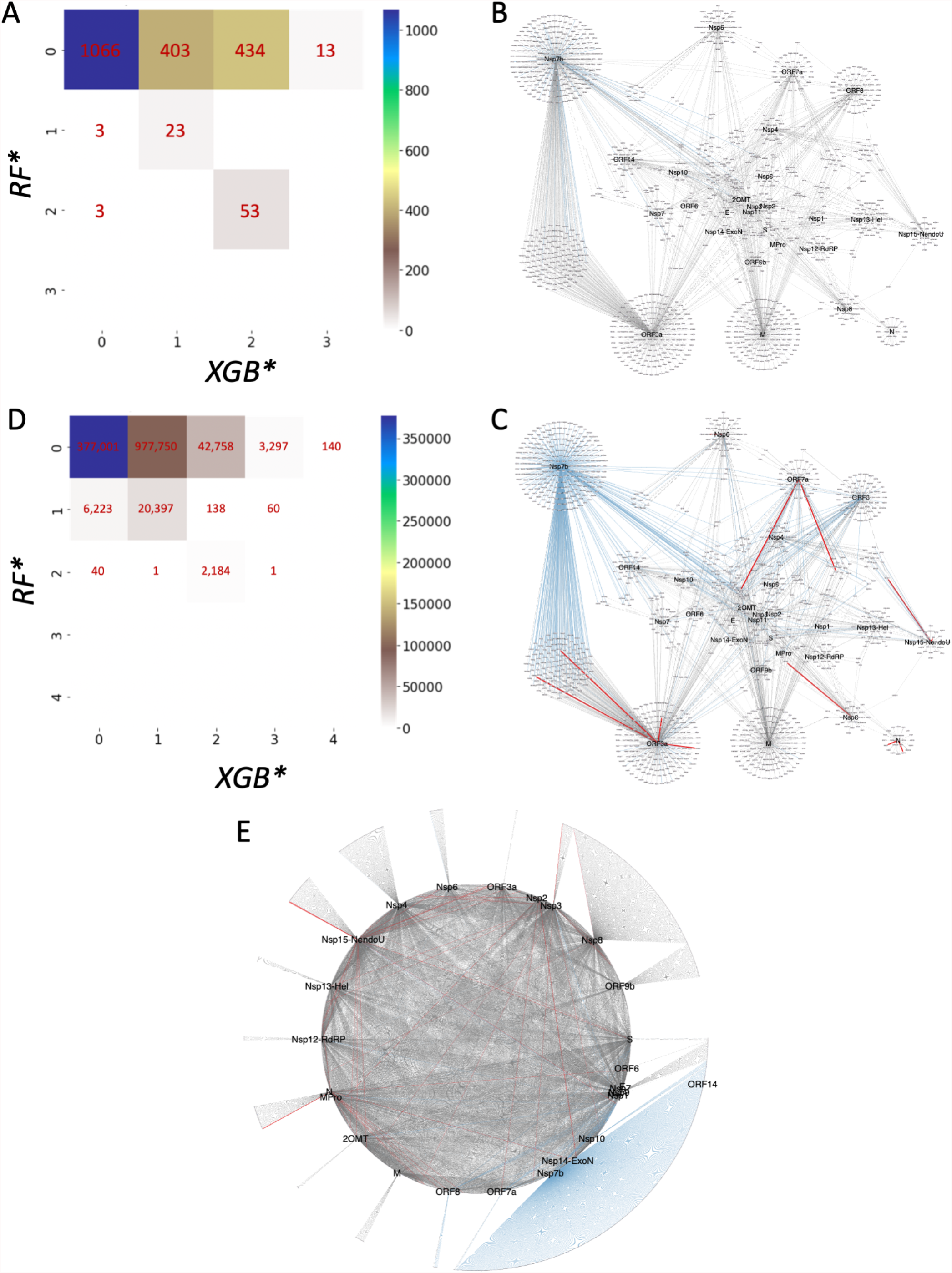
Prediction agreement between *RF\** and *XGB\**. (A) A heatmap of a confusion matrix for predictions of 1,998 PPIs from the IntAct database. (B and C) Network visualization of the 1,998 PPIs with evidence level predictions by *RF\** (B) and *XGB\** (C) based on (A). (D) A heatmap of confusion matrix for predictions of 549,990 PPIs between the SARS-CoV-2 and human proteome. (E) Network visualization of 22,781 PPIs with EC >= 1 by either *XGB\** or *RF\** from (D). Note that the EC values on the x and y axes in (A) and (D) are 0-index based. Predicted evidence levels are denoted by different edge widths and colors in (B-C) and (E): red for EC3, blue for EC2, dark grey for EC1, and light grey for EC0. Network layouts were yFiles Organic Layout for (B-C) and Circular Layout for (E) in Cytoscape.

### A virus-human interactome landscape for SARS-CoV-2

We next applied *RF\** and *XGB\** to all possible virus-human PPIs between the SARS-CoV-2 and human proteomes as described in Methods to obtain a comprehensive interactome landscape. *RF\** predicted 520,946 PPIs as EC1, 26,818 PPIs as EC2, and 2,226 PPIs as EC3 (Table S2). *XGB\** predicted 383,264 PPIs as EC1, 118,148 PPIs as EC2, 45,080 PPIs as EC3, 3,358 PPIs as EC4, and 140 PPIs as EC5 (Table S2). Prediction agreements between the two models are as follows: 377,001 PPIs for EC1, 20,397 PPIs for EC2, and 2,184 PPIs for EC3 (Fig. 4D). There are 22,781 PPIs with EC >= 2 by either *RF\** or *XGB\** (Figs. 4D and 4E).

To further support the experimental PPIs from the IntAct database analyzed above in view of the proteome-wide predictions, we compared the IntAct predictions to random subsets of the proteome-wide predictions of the same size. In other words, we performed Monte Carlo simulations for the fraction of each prediction class. With 1,000 simulations, EC3 predictions showed empirical p-value = 0 for both *RF\** (56 PPIs or 2.8%) and *XGB\** (487 PPIs or 24.4%). EC1 predictions by *RF\** (1,916 PPIs or 95.9%) showed empirical p-value = 0.016. All other predictions showed empirical p-values > 0.4.

For biological relevance of the exhaustive proteome-wide predictions, we first focus on the consensus 2,184 PPIs with EC3. Nsp7b interacts with 1,913 human proteins as the top SARS-CoV-2 protein, while two human proteins, H2AC6 and H2AC11 (histone H2A components), interact with 12 virus proteins each as the top human proteins. In fact, all of the top 15 human interacting proteins are histone H2A components. For 3,358 PPIs predicted as EC4 by *XGB\** alone, the top SARS-CoV-2 protein is Nsp3 which is predicted in 1,151 PPIs, while the top human proteins are ANKRD9 and MGAT3 which are predicted in 6 PPIs each. In the case of 140 PPIs predicted with the highest evidence of EC5 by *XGB\** alone (between 8 SARS-CoV-2 proteins and 137 human proteins) (Fig. 5A), the top SARS-CoV-2 protein is Nsp7 predicted to interact with 79 human proteins, which are significantly enriched in vascular smooth muscle contraction (CALD1, NPR2, CALML3) and Myc targets (CBX3, PES1) among others (https://maayanlab.cloud/Enrichr/enrich?dataset=9b7e9bcf2ff5a92d54529dbcbc9bbf12).

**Figure 5.**
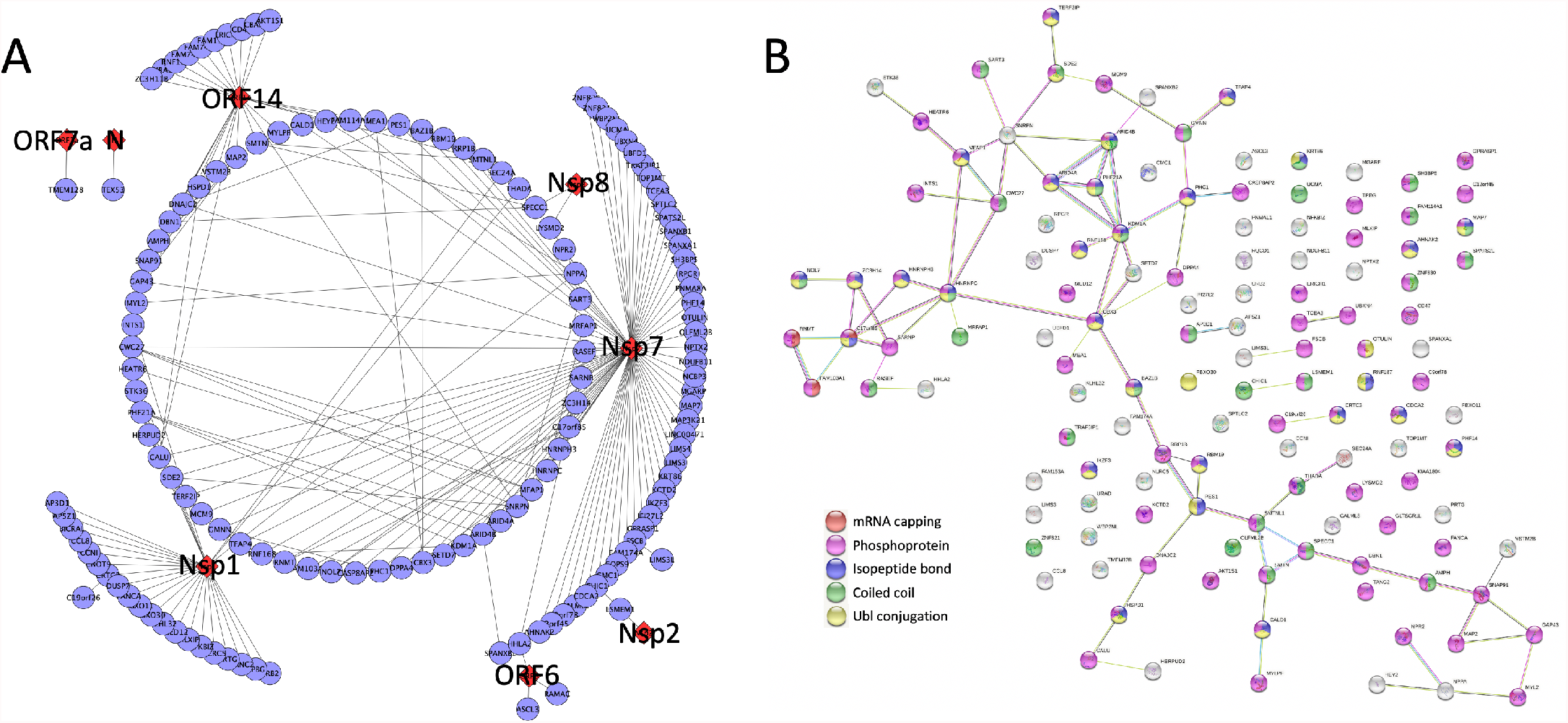
SARS-CoV-2 virus-human PPIs with the highest evidence class by *XGB\**. (A) 140 virus-human PPIs predicted for EC5 between 8 SARS-CoV-2 proteins and 137 human proteins and 81 human-human PPIs (or associations) from the STRING database. The red diamond nodes are SARS-CoV-2 proteins and the purple circle nodes are human proteins. The network was generated based on a circular layout from Cytoscape. (B) The STRING protein-protein association network of 131 human proteins from (A) and 81 associations among them. Protein nodes are color-coded by functional annotations as shown. Edge colors represent different association evidence types. Details are found in https://version-11-5.string-db.org/cgi/network?taskId=bw0zo15JMFRX.

The second most interacting SARS-CoV-2 protein is Nsp1 predicted to interact with 28 human proteins, which are significantly enriched in Hedgehog signaling pathway (STK36) and Fas signaling pathway (NPPA) among others (https://maayanlab.cloud/Enrichr/enrich?dataset=b45c89e02e13a7ed4ef51f204f94c2b2). The third most interacting SARS-CoV-2 protein is ORF14 predicted to interact with 25 human proteins, which are significantly enriched in VEGFA-VEGFR2 signaling (MYL2, AKT1S1, CALU) and LKB1 signaling (MAP2, AKT1S1) among others (https://maayanlab.cloud/Enrichr/enrich?dataset=1ef4f1d43780204d76bfeee3843cf7c7). The most frequent human interacting proteins are LYSMD2, CALU, and FAM114A1 which are predicted in 2 PPIs each. Analysis from the STRING database shows that 131 out of the 137 human proteins have 81 PPIs among themselves with an average node degree of 1.24 and a PPI enrichment p-value of 4.4e-5 (Figs. 5A and 5B) (https://version-11-5.string-db.org/cgi/network?networkId=bITYG9ekWabJ). Three proteins of mRNA capping, RNMT, FAM103A1, and C17orf85, shows the strongest association. The largest number of proteins (81) are annotated as phosphoproteins. Other proteins are annotated with Isopeptide bond (28), Coiled coil (30), or Ubl conjugation (31).

## DISCUSSION

We have developed multi-label classifiers based on RF and XGB to predict 5 evidence classes for virus-human protein-protein interactions using the Viruses.STRING database and protein sequence profiles. We computed 72 distance or similarity measures between amino acid composition profiles of interacting protein pairs as model features. *RF\** and *XGB\** achieved the mean cross-validated AUC of 67% and 74%, respectively. Two analyses of feature importance showed that cysteine and histidine are two important amino acids for virus-human PPIs, either their fractional difference or ratio. More investigation is needed to understand underlying mechanisms.

Importantly, *XGB\** showed higher prediction accuracy for experiments-based PPIs than for text mining-based PPIs, which suggests that our sequence-based features can predict physical PPIs with good accuracy. Note that the Viruses.STRING database is biased towards text mining-based PPIs, which is larger than experiments-based PPIs by more than 6-fold. Also, experiments-based PPIs are mostly in EC3, whereas text mining-based PPIs are mostly in EC1 and EC2. This means that text mining-based PPIs effectively serve as negative examples for experiments-based PPIs, which may be used for binary classification too. Hence, this allowed us to meaningfully apply our classifier to a new test set of virus-human PPIs for the new coronavirus species, SARS-CoV-2. A total of 500 SARS-CoV-2 virus-human PPIs out of 1,998 PPIs from the IntAct database has support from *XGB\** for physical PPIs with EC3 or EC4. 53 of them have additional support from *RF\** with EC3, all of which interact with Nsp7b. In addition, the fraction of EC3 predictions by either *XGB\** or *RF\** showed statistical significance by Monte Carlo simulations with respect to the exhaustive proteome-wide predictions, which gives those predictions higher priority for further investigation. Therefore, our methodology is a useful tool for predicting physical or functional virus-human PPIs, offering evidence for experimentally testable hypotheses.

Given such meaningful results, we also applied *RF\** and *XGB\** to all protein pairs between the SARS-CoV-2 and human proteomes to identify PPIs with high evidence of physical interactions (i.e., EC >= 3). There are 45,080 PPIs with EC3 and 3,498 PPIs with EC4 or EC5 predicted by *XGB\**. 2,184 of those 45,080 PPIs are also supported by *RF\** with EC3. This provides a draft landscape of the virus-human interactome for SARS-CoV-2 infection in a comprehensive and unbiased way, complementing experimentally measured PPIs. For example, *XGB\** predictions for 48,578 PPIs with EC >= 3 could greatly enhance the IntAct data of 1,998 SARS-CoV-2 virus-human PPIs. This virus-human interactome landscape could be also useful for drug repurposing studies by providing higher coverage and hence novel candidates (Morselli Gysi et al., 2021).

A limitation of our approach is lack of confidence in prediction of the low evidence classes, EC1 and EC2. They do not possess any intrinsic unique properties associated with each evidence level, unlike EC3 for experiments-based or physical PPIs. Further investigation is needed to characterize and interpret each evidence class and identify important features for each class. Alternatively, multi-class labeling might be done in different ways with different thresholds for individual evidence classes. On the other hand, one could use our tool as a binary classifier for physical PPIs as we demonstrated with EC >= 3 vs. EC < 3, or build binary classifiers based on a single threshold for combined scores. We could also benefit from data of negative controls such as decoy data. Comparative analysis with binary classifiers is beyond the scope of this study. Another limitation in this work is a subjective choice of the 72 edge features as model features. A significant model improvement might be achieved by better feature engineering for both nodes and edges.

In conclusion, our protein sequence-based multi-label classifiers are useful tools to provide different evidence or confidence levels for virus-human PPIs and applicable to virus-human interactomes for new virus species such as SARS-CoV-2.

## DATA AND CODE AVAILABILITY

Raw data are available from each of the public databases used. Full prediction results for SARS-CoV-2 are available in Supplementary Tables S1 and S2. Codes are available upon reasonable request.

## ACKNOWLEDGEMENTS

The author would like to thank Dr. Prashant Emani for helpful discussions and suggestions, Drs. Shrikant Mane and Mark Gerstein for helpful feedback and support, and the Yale Center for Research Computing for computational resources and help. This work was initiated as part of research in the author’s COVID HASTE working group at the Yale School of Engineering & Applied Science in April 2020 and partially supported by the Northeast Big Data Innovation Hub Seed Grant Program 2020.

## AUTHOR CONTRIBUTIONS

HL contributed to all aspects of the study and wrote the manuscript.

